# The double-edged effect of environmental fluctuations on evolutionary rescue

**DOI:** 10.1101/2024.08.09.607399

**Authors:** Shota Shibasaki, Masato Yamamichi

**Affiliations:** Center for Frontier Research, National Institute of Genetics, 1111 Yata, Mishima, Shizuoka 411-8540, Japan; Genetics Program, Graduate Institute for Advanced Studies, SOKENDAI (The Graduate University for Advanced Studies), 1111 Yata, Mishima, Shizuoka 411-8540, Japan; School of the Environment, The University of Queensland, St. Lucia, Brisbane, Qeensland 4072, Australia; Department of International Health and Medical Anthropology, Institute of Tropical Medicine, Nagasaki University, 1-12-4 Sakamoto, Nagasaki 852-8523, Japan; Institute for Multidisciplinary Sciences, Yokohama National University, 79-5 Tokiwadai, Hodogaya, Yokohama, Kanagawa 240-8501, Japan; Department of Ecosystem Studies, Graduate School of Agricultural and Life Sciences, The University of Tokyo, 1-1-1 Yayoi, Bunkyo-ku, Tokyo 113-8657, Japan

**Author notes:** Corresponding author: M.Y.

**Keywords:** eco-evolutionary dynamics, environmental fluctuations, evolutionary rescue, experimental evolution, green algae, mathematical model, moving optimum, quantitative genetics, rapid evolution, salt stress

## Abstract

Recent studies revealed that contemporary evolution can be rapid enough to prevent population extinction in deteriorating environments through “evolutionary rescue”. Researchers have investigated how evolutionary rescue is affected by various factors such as initial population sizes, the amount of genetic variation, and the speed of environmental changes, but few studies focused on environmental fluctuations. As the ongoing global changes are influencing the mean and variance of many environmental variables, it is crucial to explore how environmental fluctuations affect evolutionary rescue for understanding eco-evolutionary dynamics in the wild. Here we show that increasing the amplitude of environmental fluctuations around long-term deteriorating trends has negative and positive effects on evolutionary rescue by analyzing a mathematical model and conducting laboratory experiments on the green alga *Chlorella vulgaris* under increasing salinity. Increasing the amplitude of environmental fluctuations produces an episode of a huge environmental change, thereby increasing the adaptation lag between the optimal trait value and the population trait mean, which eventually causes population extinction in model simulations. On the other hand, large environmental fluctuations can increase trait variance within a population, potentially promoting adaptive evolution. Indeed, we observed that algal strains that experienced large environmental fluctuations could grow in a harsh environment whereas strains that experienced smaller or no environmental fluctuations could not. These results suggest that we will be able to promote or prohibit evolutionary rescue in nature by carefully considering the double-edged effect of environmental fluctuations on evolutionary rescue.

## 1 Introduction

Classic studies assumed that evolutionary processes are much slower than ecological processes (Darwin, 1859; Slobodkin, 1961), but recent studies revealed that evolutionary dynamics can be rapid enough to affect contemporary ecological dynamics (Thompson, 1998; Hairston et al., 2005; Carroll et al., 2007; Schoener, 2011; Reznick, 2013; Hendry, 2017). One important consequence of rapid contemporary evolution is “evolutionary rescue” where population extinction is avoided by rapidly adapting to harsh environments (Gomulkiewicz and Holt, 1995; Kinnison and Hairston, 2007; Gonzalez et al., 2013; Alexander et al., 2014; Carlson et al., 2014; Bell, 2017). By facilitating or prohibiting evolutionary rescue, we may be able to prevent the extinction of endangered species or promote the extinction of pathogenic bacteria, respectively. Evolutionary rescue has been, therefore, intensively studied due to its broad implications for conservation (Kinnison and Hairston, 2007; Alexander et al., 2014), natural resource management (Vander Wal et al., 2013), agriculture (Kreiner et al., 2018; Madgwick et al., 2024), epidemiology (Golas et al., 2021), and medicine (Alexander et al., 2014; Patil et al., 2024).

Previous studies explored the effects of various factors on evolutionary rescue by using theoretical and empirical approaches (reviewed in Alexander et al., 2014; Carlson et al., 2014; Bell, 2017). Examples include initial population sizes (Gomulkiewicz and Holt, 1995; Bell and Gonzalez, 2009; Samani and Bell, 2010; Ramsayer et al., 2013), standing genetic variation (Gomulkiewicz and Holt, 1995; Orr and Unckless, 2008; Agashe et al., 2011; Ramsayer et al., 2013; Orr and Unckless, 2014), species interactions (Osmond and de Mazancourt, 2013; Yamamichi and Miner, 2015; Osmond et al., 2017; Cortez and Yamamichi, 2019; Hermann and Becks, 2022; Morita and Yamamichi, 2023), sexual reproduction (Lachapelle and Bell, 2012; Petkovic and Colegrave, 2019, 2023; Kawaguchi and Yamamichi, 2024), genetic architecture (Orr and Unckless, 2008; Gomulkiewicz et al., 2010; Uecker and Hermisson, 2016; Yamamichi, 2022), and the speed of environmental deterioration (Perron et al., 2008; Bell and Gonzalez, 2011; Lindsey et al., 2013; Killeen et al., 2017; Liukkonen et al., 2021). However, few studies focused on the effects of environmental fluctuations on evolutionary rescue. The ongoing global changes influence the mean and variance of many environmental variables: for example, the global warming not only increases mean temperatures but also causes extreme events such as severe heat waves (Meehl and Tebaldi, 2004; Fischer et al., 2021). Previous studies demonstrated that faster environmental deterioration rates prevent evolutionary rescue in broad taxa of organisms (e.g., bacteria, green algae, and yeasts) under various stressors (e.g., antibiotics, salt, and temperature) (Perron et al., 2008; Bell and Gonzalez, 2011; Lindsey et al., 2013; Killeen et al., 2017; Liukkonen et al., 2021) by increasing the “lag load” due to the difference between the population mean trait and optimal trait in the environment (Maynard Smith, 1976). Most of these studies, however, changed the environment at constant rates (i.e., no environmental fluctuations). Therefore, it is crucial to explore how environmental fluctuations affect evolutionary rescue to understand eco-evolutionary dynamics in the wild.

In this study, we focus on the effects of short-term environmental fluctuations on evolutionary rescue when the environment shows a long-term deteriorating trend. Environmental fluctuations alone can cause population extinction by decreasing the geometric growth rate (Lewontin and Cohen, 1969; Lande, 1993; McLaughlin et al., 2002). Chevin et al. (2017) considered rapid evolution to an optimum trait value and showed that environmental fluctuations can substantially increase extinction risk by decreasing the expected growth rate and increasing the variance of population size (Marrec and Bank, 2023; Xu et al., 2023). Such “stochastic load” (Ashander et al., 2016) as well as the lag load can prevent evolutionary rescue in general when environmental fluctuations are combined with directional environmental changes (Bürger and Lynch, 1995). On the other hand, Peniston et al. (2020) showed that autocorrelated environmental variation can promote evolutionary rescue when evolutionary rescue is unlikely to occur (see also Peniston et al., 2021). In addition, environmental fluctuations can promote the maintenance of genetic variation (Ellner and Hairston, 1994; Yamamichi et al., 2023), which may promote adaptive evolution (Yamamichi et al., 2019) and prevent extinction. Compared to these theoretical studies, there are few empirical studies on environmental fluctuations in the context of evolutionary rescue. Notably, Hao et al. (2015) showed that environmental fluctuations under deteriorating trends have an evolutionary cost and an ecological benefit: the evolutionary cost arises because environmental fluctuations sometimes reverse the direction of selection pressure and increase the lag load. On the other hand, the ecological benefit occurs because when the amplitude is large enough, fluctuations can temporally improve the environmental condition in the deteriorating trend (i.e., “temporal environmental amelioration”) and may recover a population size temporally. Hao et al. (2015)’s evolutionary experiment of phages shows that the evolutionary cost outweighs the ecological benefit. It still remains unclear, however, whether environmental fluctuations impede evolutionary rescue in general as Hao et al. (2015) compared only the two scenarios without theoretical investigations.

Here, we explore how the magnitude of environmental fluctuations influences evolutionary rescue in deteriorating environments by combining theoretical and experimental approaches. By simulating a general quantitative genetic model (Lion et al., 2023), we analyzed how the amplitude of environmental fluctuations affects eco-evolutionary dynamics of a single species adapting to a deteriorating environmental trend. By conducting laboratory experiments of a unicellular freshwater green alga *Chlorella vulgaris*, we observed how the algal population adapted to increasing salinity stress for 12 weeks with (i) no variation in the elevating salinity stress, (ii) a small variation in the elevating salinity stress without reversed selection pressure (i.e., salinity always increases), and (iii) a large variation with reversed selection pressure (i.e., salinity sometimes decreases). After the 12 weeks, all three conditions reached a high salinity level (0.6M NaCl) (Fig. 1). Then, we put them to a harsher environment (1M NaCl) to see how they grew there. Our model simulations demonstrated that the larger amplitude of environmental fluctuations in general prevented evolutionary rescue as shown by previous studies (Bürger and Lynch, 1995; Ashander et al., 2016; Hao et al., 2015; Chevin et al., 2017). The simulations also showed that larger environmental fluctuations can result in larger trait variance, which may promote adaptation (i.e., an *evolutionary benefit*). Consistent with this pattern, *C. vulgaris* that experienced the large environmental fluctuations could grow in the harsh environment (1M NaCl) while the strains that experienced no or small environmental fluctuations could not (Table 1). Therefore, our results suggest that the evolutionary cost of environmental fluctuations is generally larger than the ecological benefit, but environmental fluctuations provide an evolutionary benefit by producing large trait variation. As the original (ancestral) strain quickly went extinct in the harsh environment, we concluded that the observed salinity tolerance arose via *de novo* mutations during the evolution experiment.

**Figure 1:**
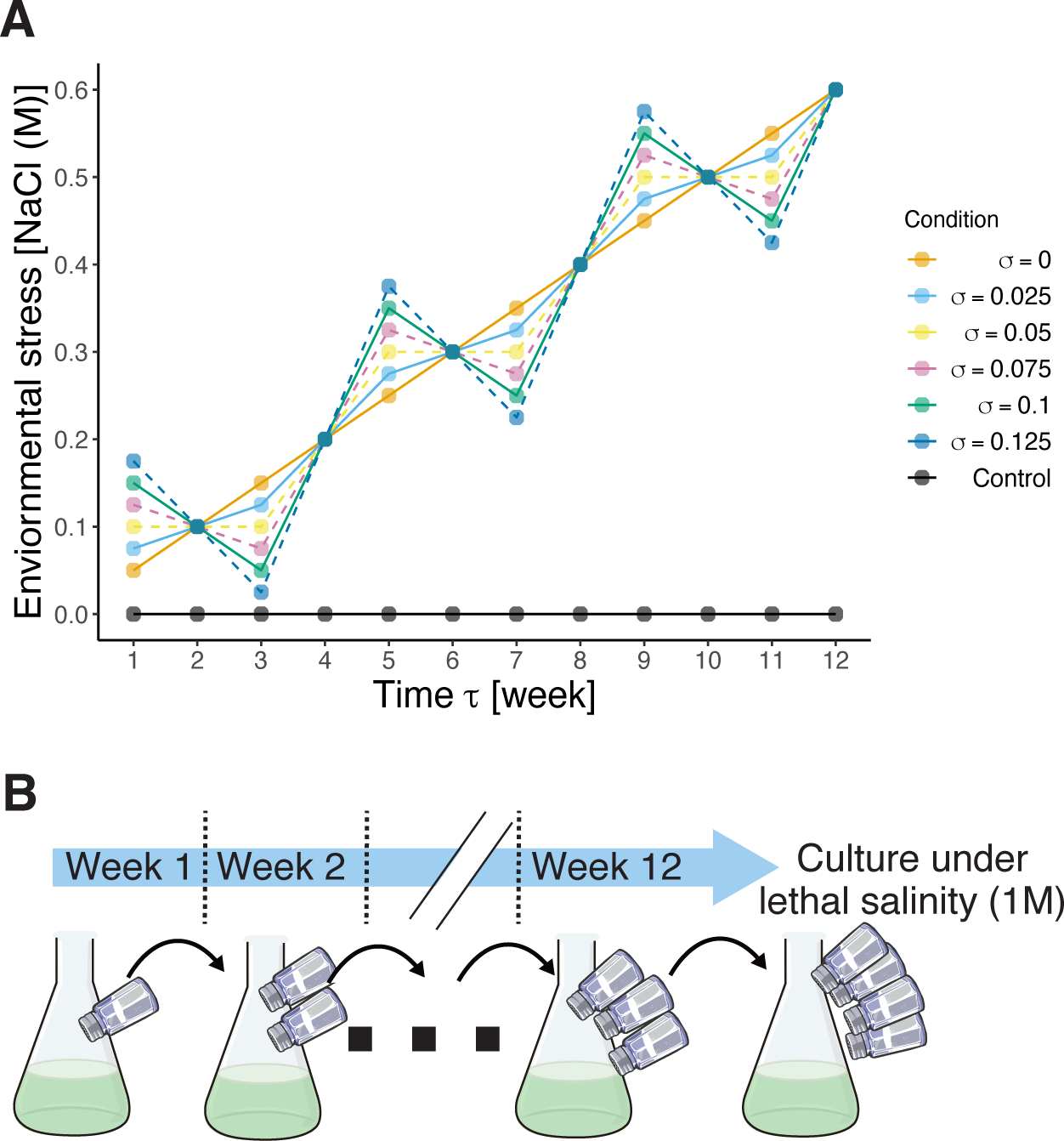
Schematic representation of the environmental changes in the simulations and experiments. A: The dynamics of environmental stress level (the salt concentration in the evolution experiment) over time. The long-term average of the environmental deterioration rate is fixed 0.05. The large amplitudes of environmental fluctuations (*σ >* 0.05), however, cause temporal environmental amelioration. The squared-brackets represent the units in the experimental evolution. The solid conditions correspond to the those in the evolution experiment. B: *C. vulgaris* was grown under varying salinity concentrations (following either control, linear: *σ* = 0.0, small-fluctuation: *σ* = 0.025, or large-fluctuation: *σ* = 0.1 conditions on panel A) for 12 weeks and then cultivated under lethal salinity stress (i.e., 1M NaCl). We transferred 1 ml of growth medium every week to a fresh medium after removing the supernatant. The salt icons by Servier https://smart.servier.com/ are licensed under CC-BY 3.0 Unported https://creativecommons.org/licenses/by/3.0/. The flask icon by DBCLS https://togotv.dbcls.jp/en/pics.html is licensed under CC-BY 4.0 Unported https://creativecommons.org/licenses/by/4.0/.

**Table 1:** Summary of the findings.

|  | Simulations | Experiments | Consistency |
| --- | --- | --- | --- |
| Likelihood of evolutionary rescue | ↘ (Fig. 2A) | No difference (Fig. 4A) |  |
| Adaptation lag (if rescued) | ↗ (Fig. 3A) | ↗ (Figs. 4B–E) | ✓ |
| Trait variance (if rescued) | ↗ (Fig. 3B) | ↗ (Fig. 5) | ✓(indirectly) |

## 2 Material and Methods

### 2.1 Mathematical model

To investigate how environmental fluctuations affect evolutionary rescue, we considered a simple eco-evolutionary model of a single population by using the theoretical framework of quantitative genetics assuming logistic growth and a normal trait distribution with small variance (Lion et al., 2023). Dynamics of the population density *N*, the mean trait *Z̅*, and the (additive) genetic variance of the trait *V* can be written as following:

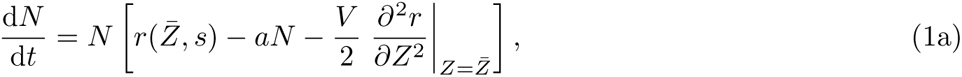

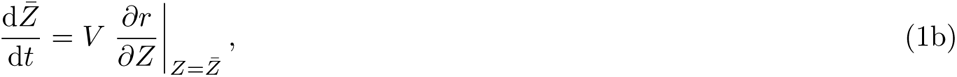

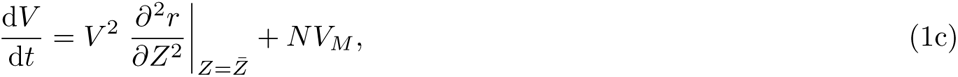

where *r*(*Z, s*) is the per-capita intrinsic growth rate, which is a function of the trait *Z* and the environmental stress level *s*, *a* is the parameter of negative density-dependence due to intraspecific competition, and *V_M_* is the coefficient for mutational variance. Population dynamics has three components: intrinsic growth for a given stress level and mean trait value (*r*), negative density-dependence (*aN*), and standing variance load due to deviations from the mean trait (*V/*2 * *∂*^2^*r/∂Z*^2^). Mean trait dynamics is determined by the two components: additive genetic variance and the fitness gradient (Lande, 1976; Iwasa et al., 1991; Taper and Case, 1992; Abrams et al., 1993). Variance dynamics is affected by selection and mutation supply. Here, we assumed that increasing the population density *N* results in a larger mutation supply.

The growth rate *r* was assumed to decrease when there is a difference between the trait mean *Z* and the environmental stress level *s* (i.e., a matching interaction). In the case of algae under salt stress, this can be because there is an optimal resistance against salinity (e.g., the osmoregulation ability), and the insufficient resistance level (*Z < s*) and the excessive resistance level with some costs (*Z > s*) can decrease the growth rate. We also assumed that high stress level decreases growth rate even when the trait mean matches the stress level because organisms may not be able to perfectly adapt to high stress. Thus, the growth function *r*(*Z, s*) can be defined as follows:

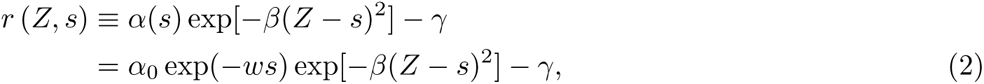

where *α*_0_ is the basic birth rate, *w* is the coefficient for the negative effect of environmental stress, *β* is the coefficient for representing the negative effect of mismatch between the trait mean and environmental stress, and *γ* is the basic mortality rate. When there is no environmental stress (*s* = 0) and the trait is zero (*Z* = 0), *α*_0_ − *γ >* 0 gives the maximum growth rate. When there is an environmental stress (*s >* 0) but the population meant trait matches with the optimal trait value (*Z* = *s*), the growth rate is *α*_0_ exp(−*ws*) − *γ*. The difference between *Z* and *s* further decreases the growth rate (i.e., lag load).

The first- and second-order derivatives of *r* are written as follows:

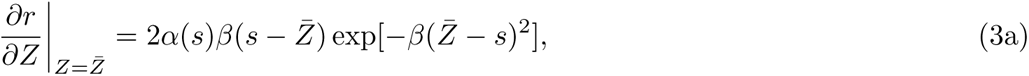

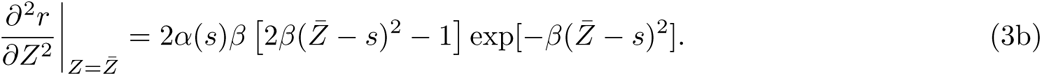

Therefore, *s − Z̅* determines the sign of the fitness gradient in the mean trait dynamics: when *Z̅ < s*, the mean trait increases whereas it decreases when *Z̅ > s*. On the other hand, 2*β*(*Z̅* − *s*)^2^ *−* 1 determines variance dynamics (see Results).

In simulations, we changed the environmental stress level every *T_f_*time step. The environmental stress level at the (*τ* + 1)th change is represented as following:

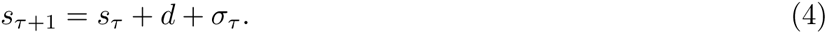

When there is no environmental fluctuation (*σ_τ_* = 0), the environmental stress steadily increases with the parameter *d >* 0. The parameter for environmental fluctuations, *σ_τ_*, is defined as follows:

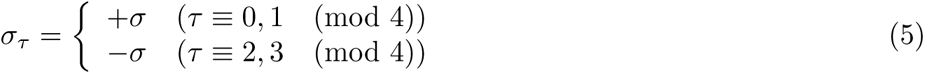

where (mod 4) returns the remainder of a division after *τ* is divided by 4. The amplitude of environmental fluctuations is determined by the parameter (*σ* ≥ 0). We set the initial environmental condition as *s*_0_ = 0 and started simulations from *τ* = 1 in this manuscript.

The dynamics in Eq. (4) has two nice characteristics. First, the long-term environmental deterioration trend is determined by *d* and is not affected by *σ*. Second, environmental fluctuations can temporarily decrease the environmental stress when *σ* is larger than *d* (i.e., temporal environmental amelioration: Hao et al., 2015). If *d > σ >* 0, on the other hand, the environment continues to deteriorate without temporal amelioration. Previous studies (Bürger and Lynch, 1995; Yamamichi et al., 2019; Peniston et al., 2021) implemented environmental fluctuations as Gaussian random variables, but the Gaussian distribution does not allow us to compare environmental deterioration with and without temporal amelioration. In addition, because we assume that the environmental stress level is zero or positive (i.e., *s* ≥ 0), the Gaussian distribution is unsuitable.

We sampled 10,000 sets of parameter values from the following distributions to compare eco-evolutionary dynamics over *σ*:

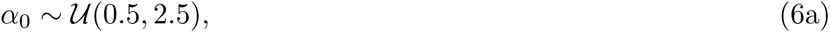

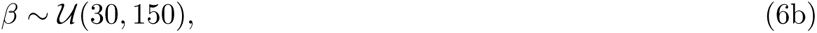

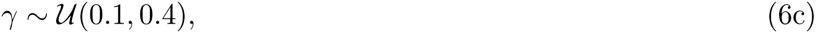

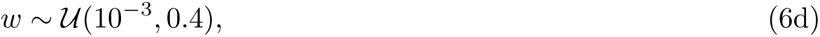

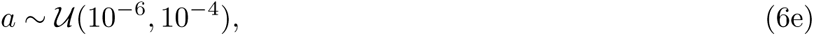

where *u*(*a, b*) represents the uniform distribution between *a* and *b*. The ranges of these parameter values were chosen so that organisms do not go extinct (i.e., the final population density in simulations is larger than a certain threshold: here, we assumed 10) when *σ* = 0. We also confirmed that all sampled parameter sets lead population extinction in the absence of evolution (i.e., the growth rates are negative when (*Z̅, s*) = (0, 0.6)). Because extinction is inevitable without evolution, population persistence in the simulations is regarded as evolutionary rescue. We fixed the values of the average environmental degradation rate (*d* = 0.05), mutation variance (*V_M_*= 10*^−^*^10^), and the frequency of the environmental changes (every *T_f_*= 50 time steps) in the main text. We continued the simulation until *τ* = 12 (600 time steps) as we did in our algal experiments (see below). We interpreted that populations did not go extinct if *N >* 10 at the end of simulations, but the simulation results were robust against the threshold (Fig. S1). For each set of parameters (*α*_0_*, β, γ, w*, and *a*), we ran simulations at *σ* = 0, 0.025, 0.05*, …,* 0.125 to compare how *σ* changed eco-evolutionary dynamics (Fig. 1).

We used the ode() function in the deSolve library version 1.38 (Karline Soetaert et al., 2010) in R version 4.3.1 (R Core Team, 2023) to simulate the eco-evolutionary dynamics. The parameter sets used in this analysis are available in supporting information.

### 2.2 Strain and growth medium

*C. vulgaris* is a unicellular and asexually-reproducing freshwater green alga widely used in laboratory experiments to investigate eco-evolutionary dynamics (Fussmann et al., 2000; Yoshida et al., 2003; Meyer et al., 2006; Kasada et al., 2014; Fisher et al., 2016). While those previous studies focused on genetic variation of the algal species along the defense-growth trade-off, here we focus on adaptation to salinity stress. Salinity stress was used for investigating evolutionary rescue for other green algae (*Chlamydomonas* and *Closterium*) in previous studies as salt imposes osmotic and oxidative stresses by disrupting the ion homeostasis and inhibits photosynthesis (Lachapelle and Bell, 2012; Petkovic and Colegrave, 2019, 2023; Kawaguchi and Yamamichi, 2024).

Growth rates of this species have been investigated under various NaCl concentrations, and previous studies showed that a 0.4M or higher concentration of NaCl significantly decreased growth rates (Alyabyev et al., 2007; Pandit et al., 2017). Alyabyev et al. (2007) further reported that *C. vulgaris* did not grow under 1M NaCl, which is consistent with our pilot experiments (Fig. S7). Although a low salt concentration (0.2M or less) facilitated growth in the previous studies (Alyabyev et al., 2007; Pandit et al., 2017), in our pilot experiments, the growth rates of *C. vulgaris* decreased over all salt concentrations from 0.0M to 0.6M (Fig. S6 and Table S4). Hence, we considered that salt acts as a stressor for *C. vulgaris* in our system.

We grew strain N-227 of *C. vulgaris*, obtained from the National Institute for Environmental Studies (NIES), Japan, in the C medium (Ichimura, 1971) for one week before starting the experiments. All cultures were kept at 23*^◦^*C under 14 : 10 light:dark cycle. The light intensity was 145 ± 5µmol m*^−^*^2^s*^−^*^1^. The original (ancestral) strain of *C. vulgaris* had a positive growth rate in the C medium when salinity was less than 0.6M NaCl (Fig. S6).

### 2.3 Measuring population size

We measured *C. vulgaris* cell density three times for every sample using a Countess II FL, an automated cell counter (Thermo Fisher Scientific Inc.), and reported the means and standard errors. The lowest threshold cell density that Countess II FL can accurately estimate is 1.00 10^5^ cells/ml. We concluded that strains went extinct when the mean cell density was lower than this threshold.

### 2.4 Evolution experiments

*C. vulgaris* was grown under increasing salinity stress for 12 weeks (Fig. 1A), except for the control strain. During the experiments, the algae were grown in 200 ml Erlenmeyer flasks with 20 ml of C medium (with or without added salt) with shaking (120 ± 2rpm). The average weekly salt increase was 0.05M in three experimental conditions so that the final salt concentration on Week 12 was 0.6M NaCl where the original strain could not grow (Fig. S6). The three experimental conditions differed in the amplitude of environmental fluctuations (Fig. 1B), which corresponds to *σ* in our simulations. In the linear condition, the salt condition increased by 0.05M weekly (i.e., no environmental fluctuations). The other two conditions had environmental fluctuations: (1) small-fluctuations with *σ* = 0.025M with the weekly increase of the salt concentration either larger (i.e., increased by 0.075M) or smaller (i.e., increased by 0.025M) than 0.05M, and (2) large-fluctuations with *σ* = 0.1M with the salt concentration either increased by 0.15M or *decreased* by 0.05M every week (i.e., temporal amelioration). Stains under the large-fluctuation condition experienced the same salinity stress as the linear conditions, but the order of the salt concentration differed between them. The total number of the strains grown in the experiments was 24 (three for the control condition, and seven for each of the three experimental conditions).

At the beginning of the evolution experiment, the initial cell density was set to 2.00 10^5^ cells/ml in each flask. Every week, we sampled 50 µl of the medium from each flask and measured cell densities three days after the inoculation and just before transferring to new fresh media. When the algae were transferred to new media, we sampled 1 ml of the growth medium from each sample. These samples were centrifuged for 10 minutes at 4*^◦^*C and 2120*×g*. After removing the supernatants, we inoculated the algae with 20ml of fresh media.

### 2.5 Assay of lethal salinity stress

After the 12-week evolution experiment, we grew the evolved *C. vulgaris* in fresh C medium containing 1M NaCl, which was lethal to the original strain (Fig. S7). The initial cell density was set at approximately 2 10^5^ cells/ml and cultured for two weeks with cell density measured daily.

### 2.6 Statistical analysis

All statistical analyses were performed with R version 4.3.1 (R Core Team, 2023). To calculate the growth rates, we fitted the data from each strain to an exponential curve using the `nls()` function in R:

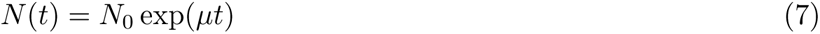

where *µ* is the growth rate (/day) and *N*_0_ is the estimated initial cell density. The initial guess of *N*_0_ in the `nls()` function was the mean initial cell density of the focal sample whereas the initial guess of *µ* was set as 0.6 in all cases.

We compared the estimated growth rates *µ* among the four scenarios at each salt concentration using the Wilcoxon rank-sum test in the `rstatix` package version 0.7.2 (Kassambara, 2023). The effect size *r* of the Wilcoxon rank-sum test was obtained from the `wilcox effsize()` function in the `rstatix` package. The *p* values were corrected by the Benjamini-Hochberg method so that the false discovery rate was 0.05 using the `p.adjust()` function in R.

## 3 Results

### 3.1 Environmental fluctuations prevented evolutionary rescue in simulations

We first investigated how the amplitude of environmental fluctuations, *σ*, affects evolutionary rescue in simulations (Eqs. (1a)-(1c)). These show that the population was less likely to persist when *σ* was larger (Fig. 2A). Extinction did not occur in more than 90% of simulations in the absence of environmental fluctuations (*σ* = 0), wheares *σ* = 0.125 decreased this fraction to less than 50%. However, this result does not exclude the possibility that the effect of *σ* depends on parameter values and larger *σ* might facilitate evolutionary rescue in some parameter space. We analyzed how often eco-evolutionary dynamics changed from extinction to survival (and vice versa) when *σ* increased by 0.025 while fixing the other parameter values. Fig. 2B shows that increasing *σ* rarely facilitated evolutionary rescue when the organisms went extinct at smaller *σ*. Meanwhile, increasing *σ* caused the extinction of organisms that were evolutionarily rescued at smaller *σ*. Figs. 2C-H show examples dynamics in our simulations. When there was no environmental fluctuation, adaptation to the moving optimum occurred (Fig. 2E) and the population did not go extinct (Fig. 2C). Note that the difference between the environmental condition and the mean trait is large when *t* = 200, and population density decreased (Fig. 2C) while variance increased (Fig. 2G). On the other hand, environmental fluctuations produced a large environmental change that resulted in an adaptation lag (Fig. 2F) and extinction (Fig. 2D). These examples clarify that larger *σ* prevented evolutionary rescue by producing a large environmental change. Overall, our simulations suggest that stronger environmental fluctuations impede evolutionary rescue.

**Figure 2:**
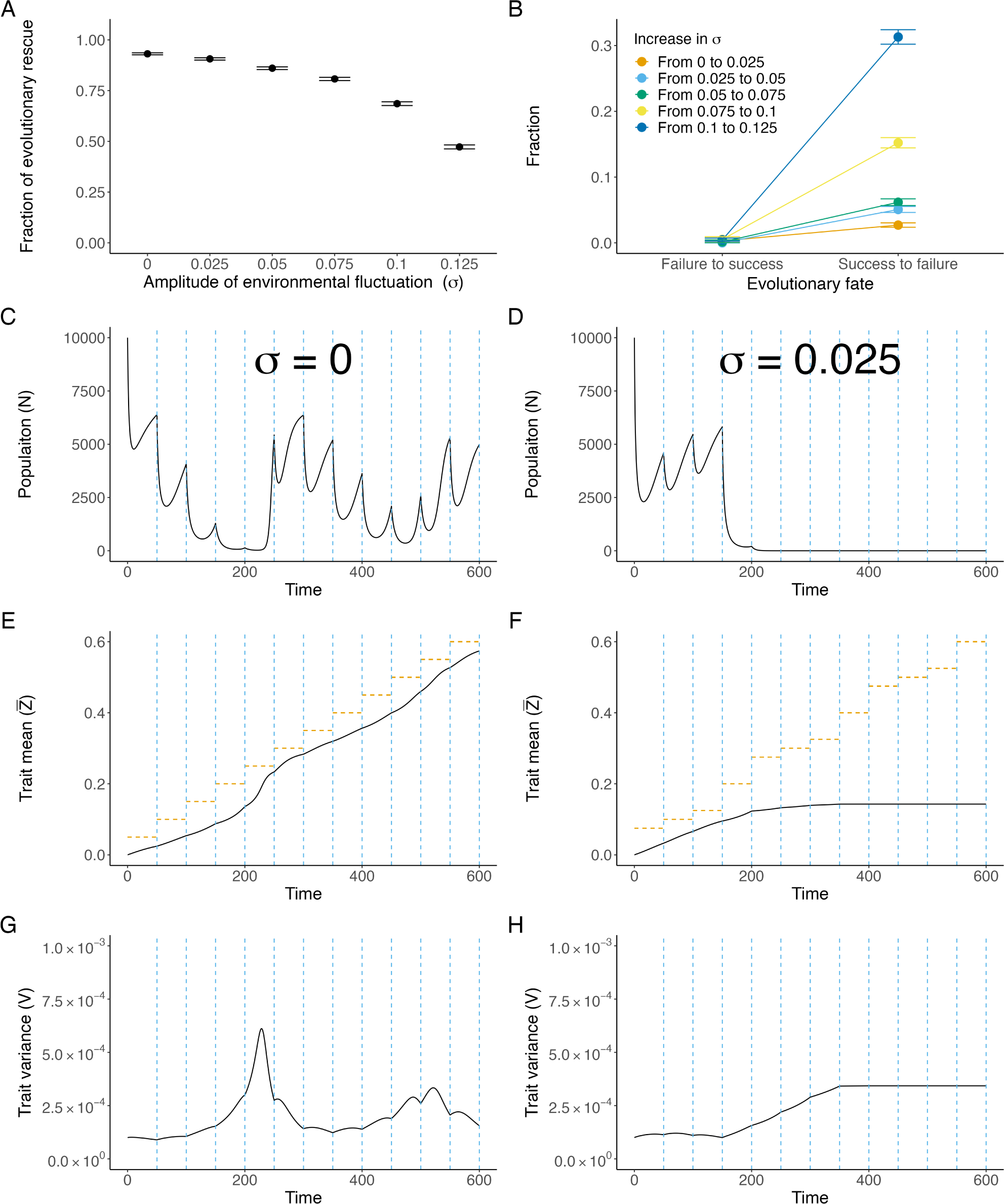
Increasing the amplitude of environmental fluctuations prevents evolutionary rescue. A: Our simulations show the fraction of successful evolutionary rescue (i.e., *N >* 10 at the end of the simulation run) decreased as the amplitude of environmental fluctuations (*σ*) increased. B: When organisms went extinct at a given *σ*, they were rarely rescued by increasing *σ*. In contrast, increasing *σ* elevated the likelihood of extinction of the rescued organisms. The error bars represent 95% HDI (highest density interval). C-H: Examples of population size dynamics (C and D), the mean trait (E and F), and its variance (G and H). The population persisted when *σ* = 0 (C, E, and G) but went extinct when *σ* = 0.025 (D, F, H). The blue vertical dashed lines indicate the time steps when the environmental stress level (horizontal orange dashed lines on panels E and F) changed. The parameter values are: *α*_0_ = 0.595194, *β* = 143.9113, *γ* = 0.190715, *w* = 0.267108, and *a* = 5.4 *×* 10*^−^*^5^.

We next analyzed how *σ* affected the mean and variance of the trait value when evolutionary rescue was successful. Fig. 3A shows that *σ* decreases the mean trait value (*Z̅*) at the end of simulations while Fig. 3B indicates that *σ* enhanced the variance (*V*) (see Table S1 for a summary of the quantile regression analysis). Figs. 3C-H show examples of the dynamics. When *σ* = 0, the mean trait steadily followed the environmental change (Fig. 3E) and the variance was almost constant over time (Fig. 3G). When *σ* = 0.125, on the other hand, the difference between the environmental condition and the trait mean was large (e.g., time steps between 0 and 50 in Fig. 3F), and the variance increased (Fig. 3H). See also Fig. S2 for the delayed evolution of the mean trait due to large *σ*. Eq. (1c) clarifies why *σ* increased *V* when there is a large difference between the environmental condition and the trait mean. Variance increases if the difference between the mean trait (*Z̅*) and the environmental stress *s_t_* is large:

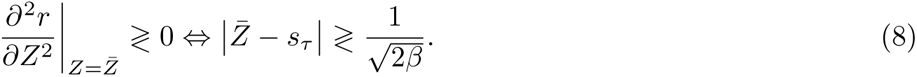

**Figure 3:**
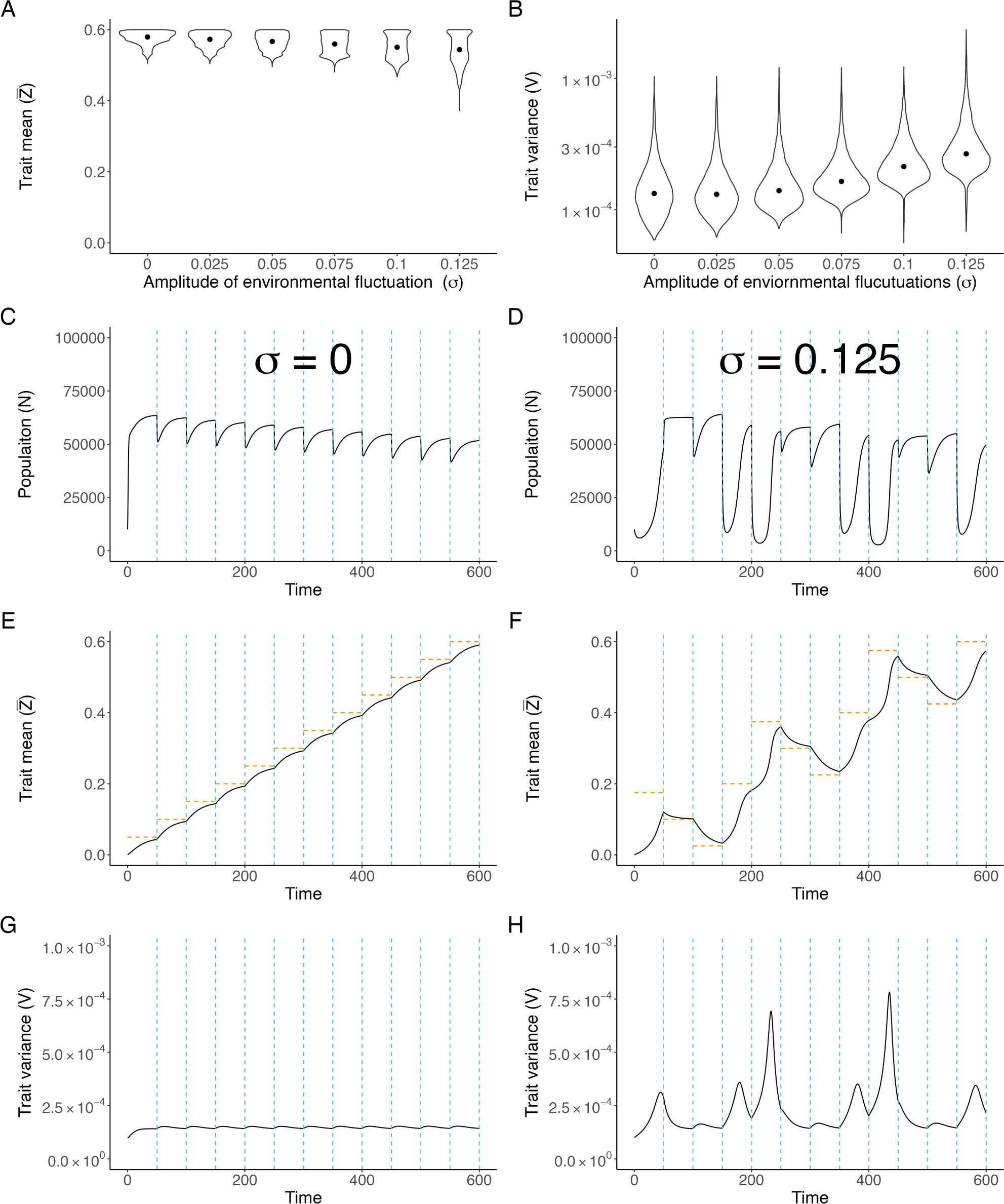
Increasing the amplitude of environmental fluctuations decreases the mean trait value but increases its variance. A and B: The distribution of mean (A) and variance (B) of the trait at the end of our simulations. While the mean trait decreased over the amplitude of the environmental fluctuations (*σ*), the variance increased. Table S1 shows the statistical details of the quantile regression. The black dots represent the median values. C-H: Examples of dynamics of the population size (C and D), the mean trait (E and F), and its variance (G and H) with no (*σ* = 0: panels C, E, and G), and strong (*σ* = 0.125: panels D, F, H) environmental fluctuations. The blue vertical dashed lines indicate the time steps when the environmental stress level (horizontal orange dashed lines on panels E and F) changed. The parameter values are as follows: *α*_0_ = 2.429814, *β* = 67.60141, *γ* = 0.116249, *w* = 0.2348825, and *a* = 3.6 *×* 10*^−^*^5^.

Larger *σ* is more likely to increase the difference between the trait and the environmental optimum because the maximum increase of the environmental stress level is *d* + *σ* in our simulations. The above inequality also indicates that the variance decreases when *Z̅* is close to *s_τ_*for decreasing the standing variance load. Indeed, Fig. 3H shows that the variance decreased when the difference is small (Fig. 3F). Thus our model simulations suggest that environmental fluctuations increase the adaptation lag and promote extinction, but they can also increase trait variance (Table 1).

### 3.2 Large environmental fluctuations delayed resistance evolution in experiments

In our experiments with *Chlorella vulgaris*, all 24 strains were maintained during the 12 weeks and there was no extinction (Fig. 4A). The strains had approximately 50–60 generations during the evolution experiments (Fig. S5); i.e., the salinity stress changed every 4–5 generations. The growth rates in the control condition were ca. 0.38/day (Fig. 4B), and they repeated similar dynamics every week (see also Fig. S8 for growth rates over the 12 weeks). The strains in the three experimental conditions (i.e., linear, small-fluctuation, and large-fluctuation conditions) had gradually decreasing cell densities due to the increasing salinity stress, but all strains continued to grow (no extinction) during the experiments. Remarkably, these strains could grow at 0.6M NaCl, where two-thirds of the original strains could not grow in our pilot experiments (Fig. S6). Assuming that at least the population size was 20 × 10^5^ cells and the number of generations was 40, each line might have 80,000,000 cells during the 12-week experiments. Because the genome size of *Chlorella vulgaris* is ca. 40Mbp (Cecchin et al., 2019) and the estimated *de novo* mutation rate of a green alga, *Chlamydomonas reinhardtii*, is ca. 10*^−^*^9^/site/generation (Ness et al., 2015), each line might have 3,200,000 mutations, which might affect salinity tolerance.

**Figure 4:**
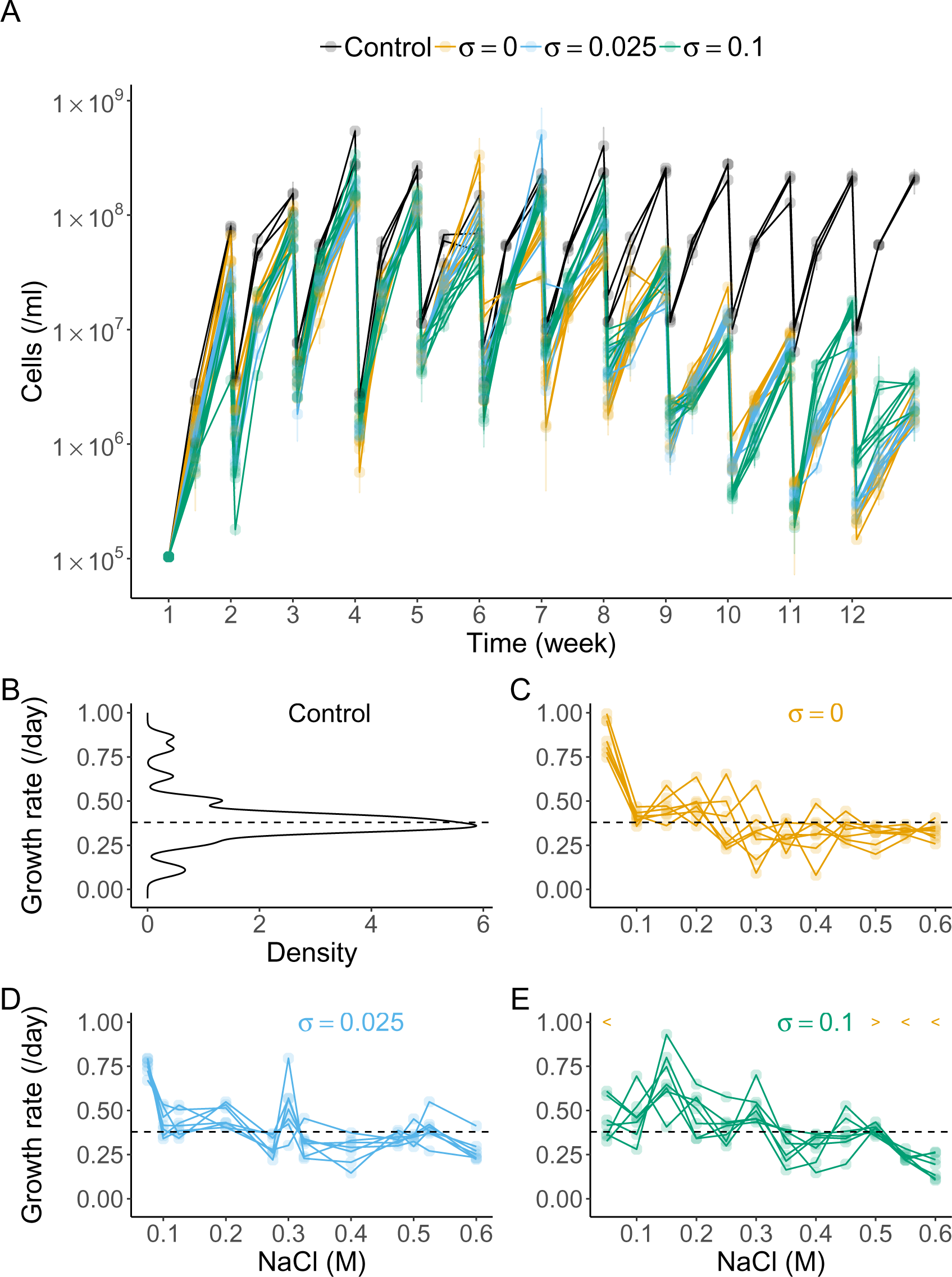
Population dynamics and growth rates during the experimental evolution. A: Population dynamics of 24 strains during the evolution experiments (black: control, orange: linear, blue: small-fluctuation, and green: large-fluctuation). Each dot represents the mean cell density of the three measurements, and each error bar shows the standard error of the three measurements. B: the distribution of the growth rates of the control condition. The dashed lines represent the median growth rates of the control condition (*µ* = 0.38). C - E: The growth rates in the three experimental conditions over the salt concentration. The orange *<* (*>*) symbols on the panel E indicate that the growth rates in the linear condition were higher (lower) than the large-fluctuation at each salt concentration. See Table S2 for the statistical details.

When we compared the growth rates in the three experimental conditions, the growth rates in the small-fluctuation condition (Fig. 4D) did not significantly differ from those in either the linear (Fig. 4C) or large-fluctuation conditions (Fig. 4E). The growth rates of the large-fluctuation condition at high salt concentration (0.55M and 0.6M), however, were smaller than those of the linear condition (one-sided Wilcoxon rank-sum test with the false discovery rate correction: *p* = 0.01, effect size *r* = 0.54 at 0.55M; *p* = 0.01, *r* = 0.77 at 0.6M: Table S2). These results were consistent with our simulations showing delayed trait evolution when *σ* was large (Fig. S2). Hence, the large environmental fluctuations delayed the evolution of resistance to the salinity stress (Table 1).

### 3.3 Large environmental fluctuations promoted growth under the lethal salinity stress

After the 12-week evolution experiment where the salinity level was increased from 0M NaCl to 0.6M NaCl, the evolved strains were exposed to the lethal salt concentration of 1M NaCl. As expected, the original strains (Fig. S7) as well as the three strains from the control condition (Fig. 5B) quickly went extinct. In contrast, the strains that experienced increasing salinity did not go extinct in this harsh environment. These strains had significantly higher growth rates than the control strains (one-sided Wilcoxon rank-sum test with the false discovery correction, Table S3). Interestingly, the strains that experienced the large-fluctuation condition grew significantly faster than the strains from the linear and small-fluctuation conditions (Fig. 5A). The seven strains from the large-fluctuation condition clearly showed positive growth rates (Fig. 5E) whereas most strains in the linear and small-fluctuation conditions persisted but did not grow (Fig. 5C and D). These results indicate that the large environmental fluctuation facilitated evolutionary rescue under lethal salinity stress.

**Figure 5:**
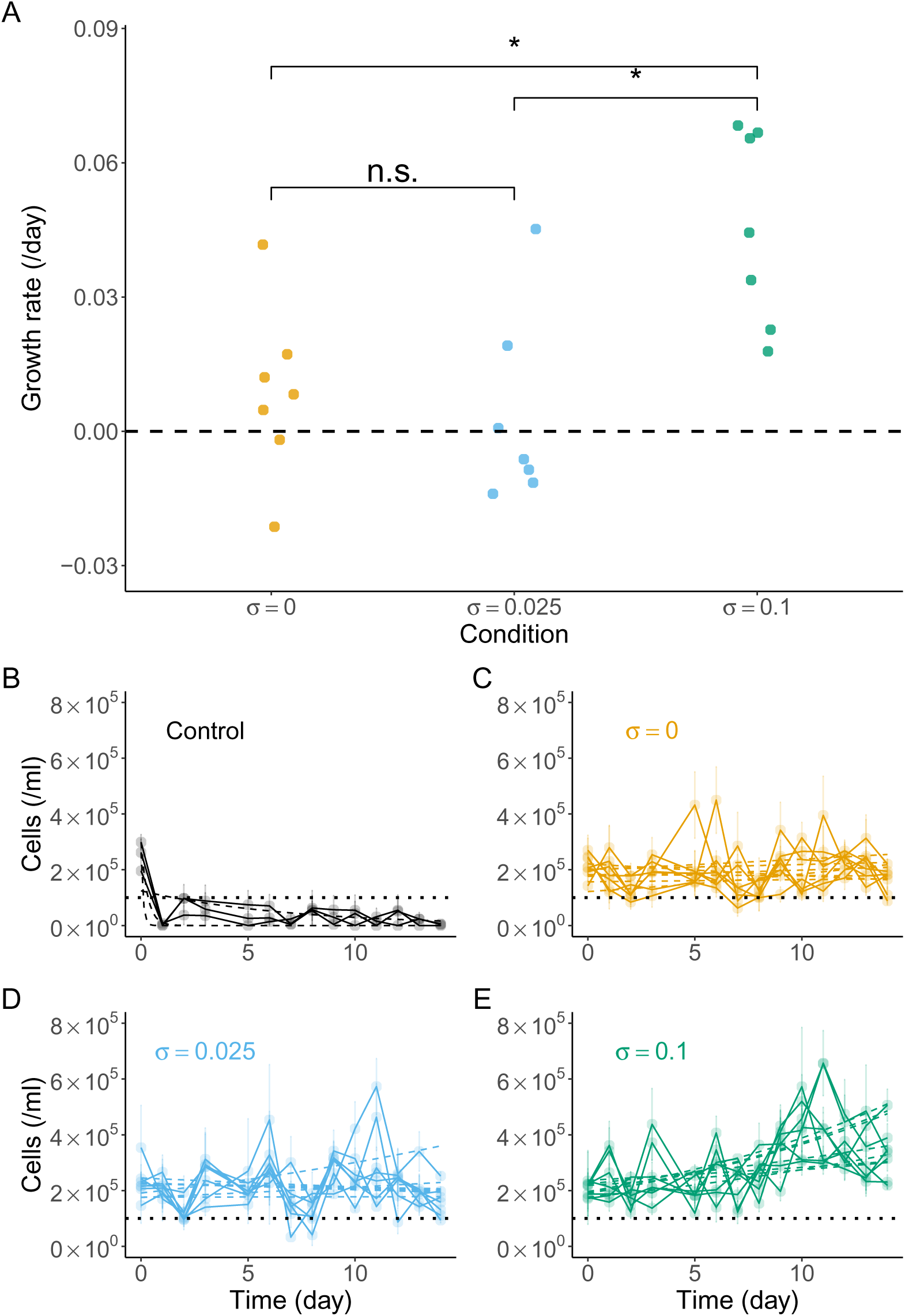
Population dynamics of evolved strains under 1M NaCl. A: The growth of the evolved strains in 1M NaCl. The asterisks represent statistical difference in the growth between the two conditions, while n.s. indicate no significance (see Table S3 for the p-values and effect sizes). The growth rates for strains of the control condition are omitted because they were very small (and negative). The growth rates of the three experimental conditions were significantly higher than the control condition. B-E: The population dynamics of each evolved strain in 1M NaCl. The dots and error bars represent the mean and the standard error of the three measurements, respectively. The dashed lines show the fitted exponential growth curves, and the dotted lines indicate the threshold for extinction.

## 4 Discussion

Environmental fluctuations affect various ecological and evolutionary processes such as geometric population growth rates (Lewontin and Cohen, 1969), population persistence (Lande, 1993), predator–prey population cycles (Vasseur and Fox, 2009), the maintenance of genetic and species diversity (Ellner and Hairston, 1994; Yamamichi et al., 2023), the rate of adaptive evolution (Yamamichi et al., 2019), and evolutionary rescue. Previous studies demonstrated that the lag load due to the long-term environmental deterioration (Perron et al., 2008; Bell and Gonzalez, 2011; Lindsey et al., 2013; Killeen et al., 2017; Liukkonen et al., 2021) and the stochastic load due to short-term environmental fluctuations (Chevin et al., 2017; Ashander et al., 2016; Marrec and Bank, 2023; Xu et al., 2023) can simultaneously impede evolutionary rescue. Here, we combine model simulations and evolutionary experiments using green algae, *Chlorella vulgaris*, to investigate how environmental fluctuations affect evolutionary rescue under deteriorating trends. We found a double-edged effect of environmental fluctuations on evolutionary rescue: larger fluctuations increase the lag between the optimal and mean trait values, which potentially results in population extinction, but also enhance evolutionary rescue by increasing trait variation (Table 1). While Hao et al. (2015) pointed out that large environmental fluctuations produce an ecological benefit by temporarily increasing the population density due to the temporarily improved environmental condition (i.e., temporal environmental amelioration), the evolutionary benefit of environmental fluctuations (i.e., enhancing trait variance) has not been discussed in previous studies.

Our simulations showed that evolutionary rescue was prevented by large amplitude environmental fluctuations, *σ*, which generalizes the findings of Hao et al. (2015). Fig. 2A clearly shows that environmental fluctuations impede evolutionary rescue even when the amplitude of environmental fluctuations is not large enough to cause temporal environmental amelioration (i.e., without reversed selection pressure) whereas Hao et al. (2015) compared evolution of phases under two conditions: without environmental fluctuations and with environmental fluctuations large enough to cause environmental amelioration. Our results are consistent with simulations in previous studies (Bürger and Lynch, 1995; Peniston et al., 2021) which showed that faster long-term environmental deterioration and larger amplitudes of environmental fluctuations increase the risk of extinction. Environmental fluctuations prevent evolutionary rescue because larger environmental fluctuations produce episodes of huge environmental change which increase the adaptation lag between the optimal and mean trait values (Figs. 3A and S2). The adaptation lag was also observed in our evolution experiments with *C. vulgaris*; the algal strains that evolved under the large-fluctuation condition grew more slowly than those without environmental fluctuations (the linear condition) in 0.55 and 0.6M NaCl (Fig. 4). However, no strains went extinct in the three-month experiment (Fig. 4A) while our simulation result shows that the likelihood of evolutionary rescue is a decreasing function of the amplitude of environmental fluctuations (Fig. 2A). This mismatch may be because the adaptation lag of the strains from the large-fluctuation condition was not large enough to cause extinction. Indeed, our simulations show that population does not always go extinct when the adaptation lag exists (Fig. 3A). Nevertheless, the strains might have gone extinct if the experiment had continued longer because a negative trend of growth rates at high salinity (0.5M or higher) in Fig. 4E suggests the increasing adaptation lag in the large-fluctuation condition. Environmental fluctuations, therefore, produce an evolutionary cost as organisms must follow the steep increase in environmental stress.

Environmental fluctuations also produce an ecological benefit (Hao et al., 2015): environmental fluctuations can maintain larger population sizes (Fig. 4A). Although Hao et al. (2015) argued that this is the benefit of environmental amelioration (i.e., temporal decrease in environmental stress), population size can recover even when the amplitude of environmental fluctuations is so small that environmental amelioration is absent (e.g., compare Figs. 2C and D between time step 150 and 200). Since larger population sizes increase the chance of evolutionary rescue, one may wonder why such a positive impact of environmental fluctuations was not observed. Our simulations suggest that this is because the ecological benefit and the evolutionary cost are two sides of the same coin. Previous studies showed that initial population sizes facilitated evolutionary rescue by increasing population sizes 7.5 times (Ramsayer et al., 2013) or by an order of magnitude (Bell and Gonzalez, 2009; Samani and Bell, 2010). To cause such a large difference in population size with and without environmental fluctuations, organisms need to be sensitive to the environments when the amplitude of environmental fluctuation is large. Mathematically, this implies that *β*, *w*, and *σ* in Eqs. (2) and (4) are large. Such conditions, however, imply that organisms are likely to go extinct because their growth rates rapidly decline when the environmental stress level steeply increases due to environmental fluctuations. In other words, the ecological benefit positively correlates with the evolutionary cost so that the benefit is unlikely to outweigh the cost.

We found, however, an *evolutionary* benefit of environmental fluctuations on evolutionary rescue. Our simulations show that trait variance increased as a function of the amplitude of environmental fluctuation (Fig. 3B). This is because the variance increases only when there is a large difference between the environmental optimum and mean trait value (see inequality (8)). Larger variance allows organisms to adapt to new environments, and this is another key factor of evolutionary rescue (Gomulkiewicz and Holt, 1995; Barrett and Schluter, 2008; Orr and Unckless, 2008; Agashe et al., 2011; Bell, 2013; Lachapelle and Bell, 2012; Orr and Unckless, 2014; Petkovic and Colegrave, 2019, 2023). Consistently, our evolved *C. vulgaris* that experienced large environmental fluctuations grew in 1M NaCl, while strains from other conditions either went extinct (the original strains and those in the control condition) or persisted but did not grow (in the linear and small-fluctuation conditions). Although we did not analyze the genome data of *C. vulgaris*, it is more likely that the resistance to salt was obtained by *de novo* mutation during the evolution experiment than standing genetic variance of the original strain because the original strain or strains from the control condition did not persist in 1M NaCl (Figs. 5B and S7). In sum, strains that experienced environmental fluctuations gain an evolutionary benefit that allows them to grow in the harsher environment, probably by increasing the trait variance.

Our quantitative genetic model was originally formulated to describe eco-evolutionary dynamics of sexually reproducing populations (Lion et al., 2023), and the assumptions are not consistent with our laboratory populations of green algae that reproduce asexually. Thus, it will be interesting in future studies to construct an individual-based model or population dynamic model with many asexual clones. In this context, consideration of the evolution of high mutation rates instead of large trait variance is an important avenue to be explored. Previous studies showed that mutation rates can evolve rapidly (Wei et al., 2022), and large environmental fluctuations might facilitate evolution of high mutation rates. To understand the role of mutation rates, it will also be important to explore *Chlorella* genome. Since *C. vulgaris* is a model organisms, genes related to salinity stress tolerance have been identified (Abdellaoui et al., 2019). Future studies can investigate which genetic changes were responsible for the experimental evolution we observed and whether genetic variance increased in the large-fluctuation condition or not. It should be noted that we did not provide direct evidence whether phenotypic variance was larger in the large-fluctuation condition than the other conditions. Although large fluctuations increased the trait variance in simulations, it was transient; once the mean resistance level became close enough to the environmental optimum, the variance started to decrease (see inequality (8)). Qualitative results of the simulations depend on parameters that affect the rate of evolution of the mean resistance level; e.g., frequency of environmental changes *T_f_*, and mutational variance *V_M_*(Figs. S3, and S4). Nevertheless, our simulations from the broad parameter ranges suggest the robust prevention of evolutionary rescue and increase of the trait variance over the amplitude of environmental fluctuations (Figs. 2A and 3B; see also Figs. S3, and S4). Therefore, we encourage further studies investigating how environmental fluctuations affect rapid evolution and evolutionary rescue.

In conclusion, our mathematical model simulations and experimental evolution of green algae suggest that environmental fluctuations have a double-edged effect on evolutionary rescue under deteriorating environments. On the one hand, introducing environmental fluctuations delay trait evolution to the moving optiumum, which can cause extinction. On the other hand, large fluctuations can increase the trait variance, allowing the organisms to grow in harsh environments.

## Data availability

Codes and data used in this manuscript are available on the following Github repository https://github.com/ShotaSHIBASAKI/The_double-edged_effect_enviornmental_flucutuation_ER.

## Acknowledgement

*C. vulgaris* (NIES-227) was provided by NIES through the National BioResource Project of the Ministry of Education, Culture, Sports, Science and Technology (MEXT), Japan. We thank Mayumi Aono for helping the experiments as a technician. We are grateful to Drs. Nelson G. Hairston Jr. and Andrew D. Letten for their fruitful comments on the earlier version of this manuscript. This study was supported by the Center for Frontier Research at the National Institute of Genetics, Japan Society for the Promotion of Science (JSPS) Grant-in-Aid for Scientific Research (KAKENHI) Grant Numbers JP19K16223, JP20KK0169, JP21H02560, JP22H02688, and JP22H04983, Japan Science and Technology Agency (JST) Core Research for Evolutional Science and Technology (CREST) JPMJCR23N5, Inamori Research Grant, and Australian Research Council (ARC) Discovery Project DP220102040 to MY.

## Author contribution

S.S. and M.Y conceptualized the project, S.S. conducted the experiments, analyzed the model, performed the analysis, and wrote the original draft. S.S. and M.Y. revised and edited the manuscript. M.Y. supervised the project and acquired the funding.

## SI 1 Supplementary Figures and Tables

**Figure S1:**
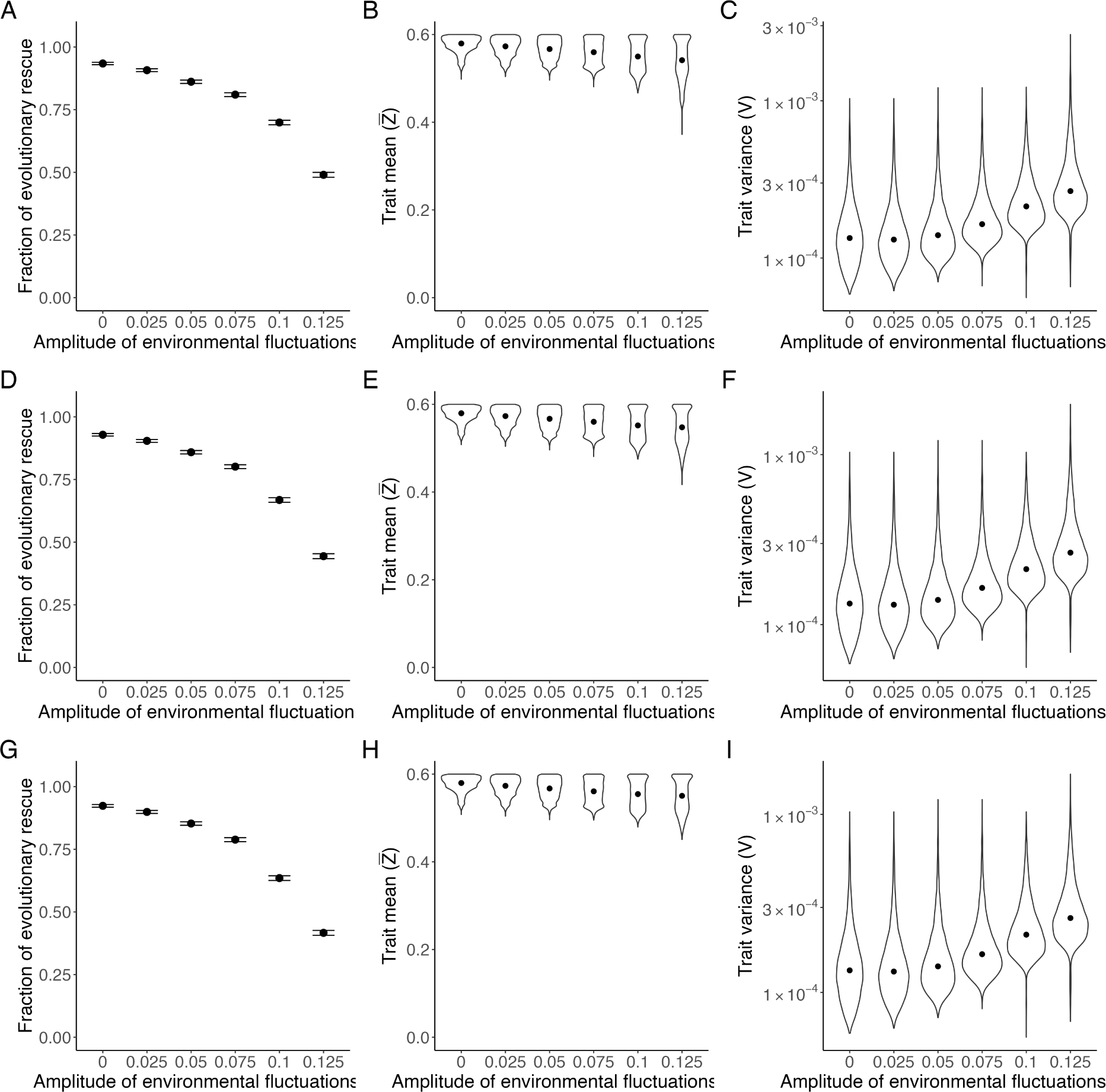
Sensitivity analysis against the threshold of extinction. The fraction of successful evolutionary rescue (left), the distribution of the mean trait when evolutionary rescue succeeded (center), and the distribution of variance of the trait when evolutionary rescue (right) over the amplitude the environmental fluctuations. The threshold of extinction was *N >* 1 (top), *N >* 100 (middle), and *N >* 1000 (bottom), respectively.

**Figure S2:**
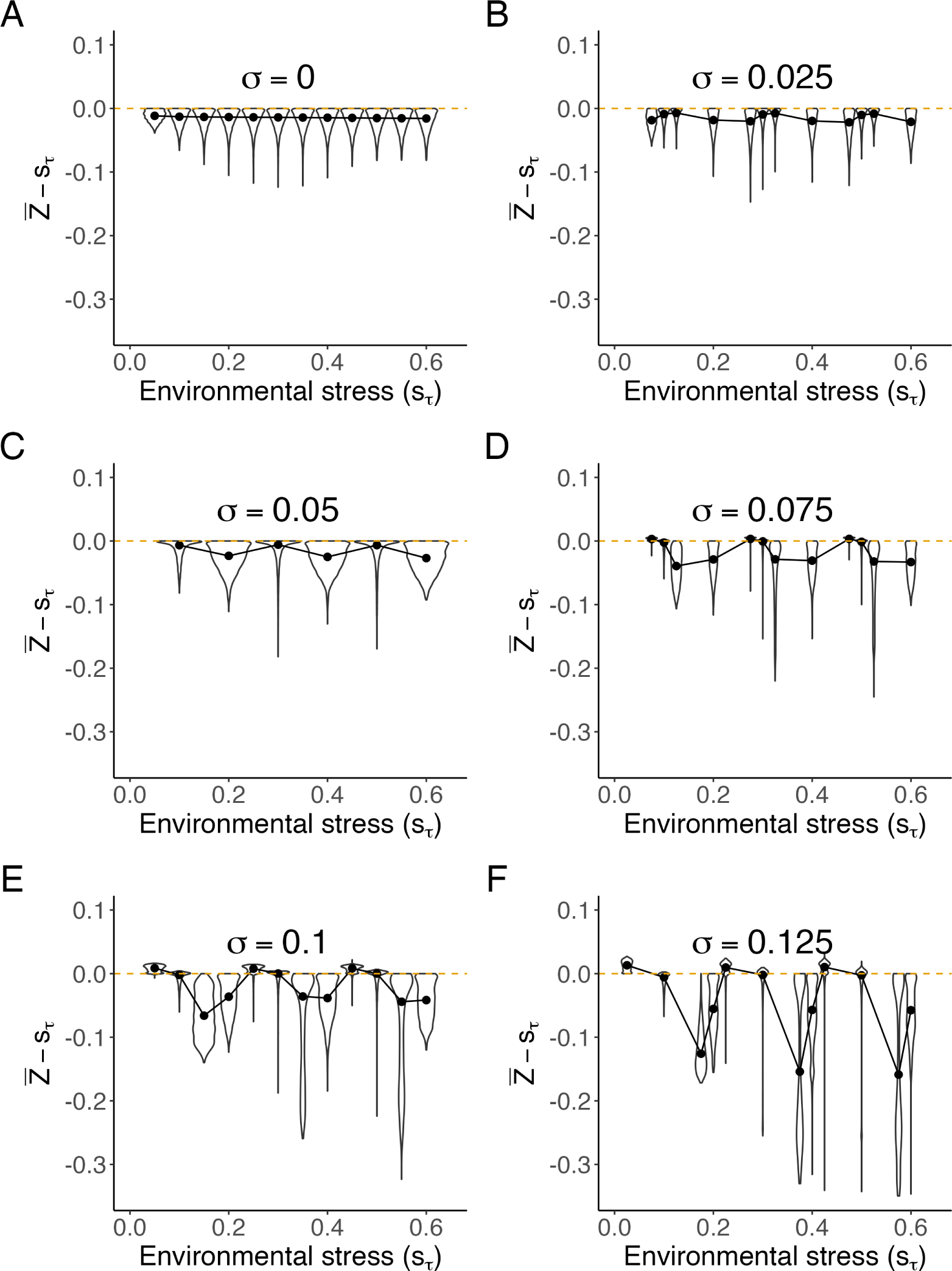
Difference between the mean trait and environmental optimum in the simulations. Difference between the mean trait (*Z̅*) just before the environmental change and the environmental optimum (*s_τ_*) is shown over the environmental stress level. The orange dashed line represent *Z̅* = *s_τ_* (i.e., optimal trait). The black dots represent the median values. To avoid the effects of different number of samples, we plotted the simulation data whose parameter value allowed the evolutionary rescue at all *σ* (in total 4,711 simulation data for each panel).

**Figure S3:**
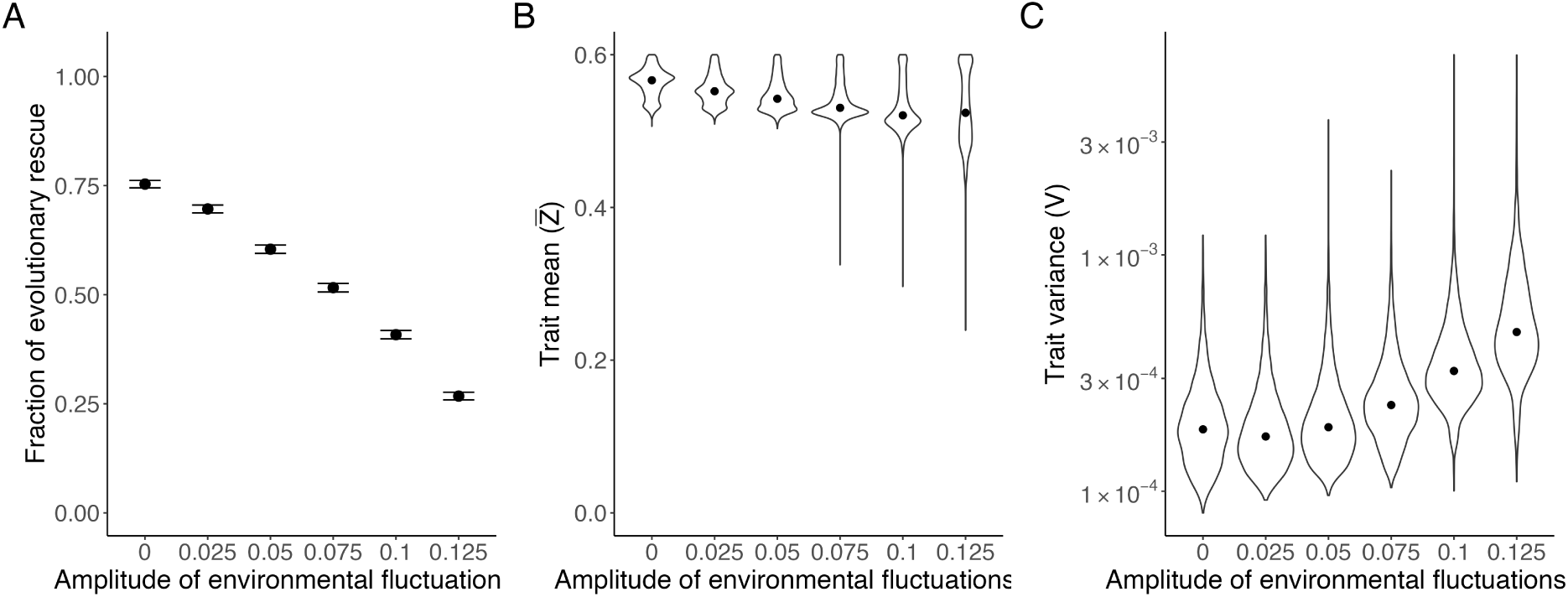
Simulation results when *T_F_* is short. We increased the frequency of environmental change by setting *T_F_* = 25 and performed 10,000 simulations as in the main text. A: The probability of successful evolutionary rescue (*N >* 10 at the end of simulations) over the amplitude of the environmental fluctuations. B: The distribution of mean trait when evolutionary rescue succeeded. C: The distribution of the trait variance when evolutionary rescue succeeded.

**Figure S4:**
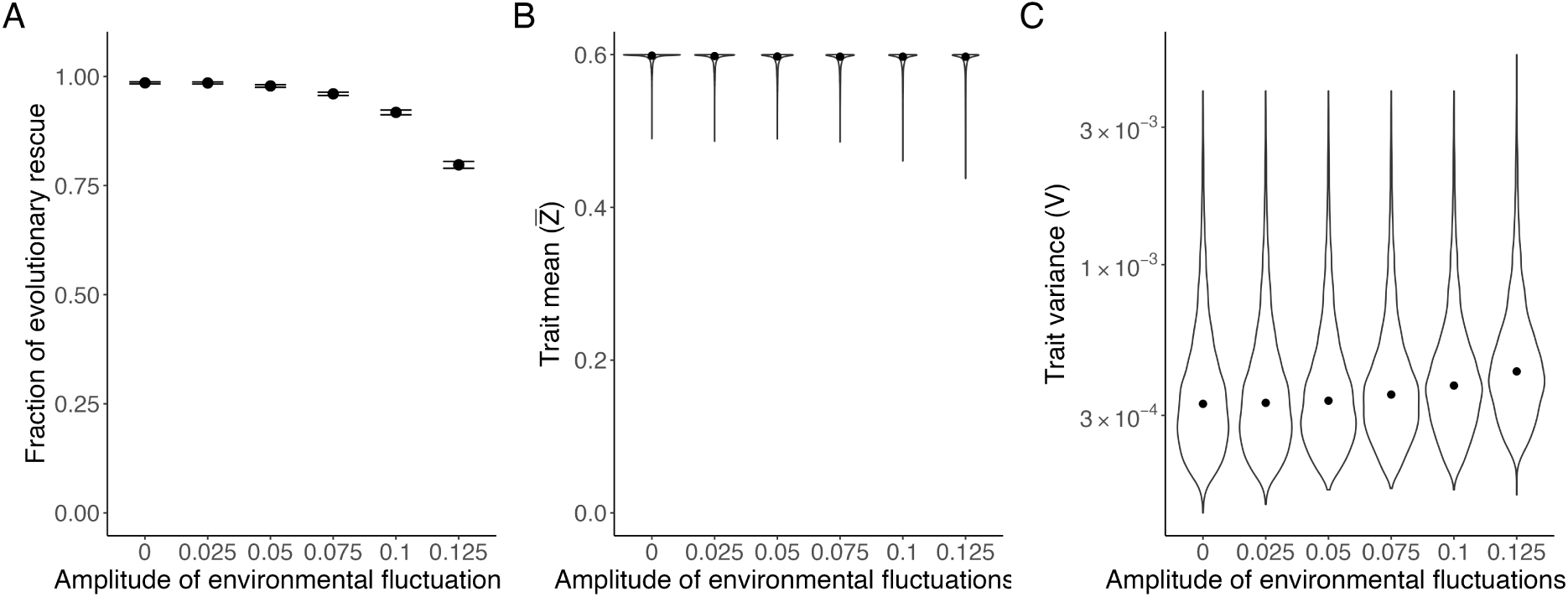
Simulation results when *V_M_* is large. We increased the mutational variance *V_M_* = 10*^−^*^9^ and performed 10,000 simulations as in the main text. A: The fraction of successful evolutionary rescue (*N >* 10 at the end of simulations) over the amplitude of environmental fluctuations. B: The distribution of mean trait when evolutionary rescue succeeded. C: The distribution of the trait variance when evolutionary rescue succeeded.

**Table S1:** The amplitude of environmental fluctuations affected mean and variance of trait values.

| Model | Coefficient | Value | Standard Error | Significance |
| --- | --- | --- | --- | --- |
| $\bar{Z} \sim \sigma$ | Intercept | 0.58002 | 0.00022 | ✓ |
| | $\sigma$ | -0.28083 | 0.00499 | ✓ |
| $V \sim \sigma$ | Intercept | 0.00011 | 0.00000 | ✓ |
| | $\sigma$ | 0.00091 | 0.00001 | ✓ |

**Table S2:** Summary of one-sided Wilcoxon rank sum tests on the growth rates in the experimental evolution.

| Pair and alternative hypothesis | Salinity | $W$ | P value | Significance | Effect size ($r$) |
| --- | --- | --- | --- | --- | --- |
| Linear > SD100 | 0.05 | 49.00 | 0.01 | ✓ | 0.84 |
| Linear < SD100 | 0.10 | 17.00 | 0.77 |  | 0.26 |
| Linear < SD100 | 0.15 | 6.00 | 0.07 |  | 0.63 |
| Linear < SD100 | 0.20 | 22.00 | 0.98 |  | 0.09 |
| Linear < SD100 | 0.25 | 18.00 | 0.78 |  | 0.22 |
| Linear < SD100 | 0.30 | 6.00 | 0.07 |  | 0.63 |
| Linear < SD100 | 0.35 | 25.00 | 1.00 |  | 0.02 |
| Linear > SD100 | 0.40 | 30.00 | 0.80 |  | 0.19 |
| Linear < SD100 | 0.45 | 18.00 | 0.78 |  | 0.22 |
| Linear < SD100 | 0.50 | 1.00 | 0.01 | ✓ | 0.80 |
| Linear > SD100 | 0.55 | 49.00 | 0.01 | ✓ | 0.84 |
| Linear > SD100 | 0.60 | 47.00 | 0.01 | ✓ | 0.77 |
| Linear < SD025 | 0.10 | 23.00 | 0.98 |  | 0.05 |
| Linear > SD025 | 0.20 | 23.00 | 0.98 |  | 0.05 |
| Linear < SD025 | 0.30 | 9.00 | 0.14 |  | 0.53 |
| Linear > SD025 | 0.40 | 30.00 | 0.80 |  | 0.19 |
| Linear < SD025 | 0.50 | 12.00 | 0.28 |  | 0.43 |
| Linear > SD025 | 0.60 | 39.00 | 0.17 |  | 0.50 |
| SD025 < SD100 | 0.10 | 23.00 | 0.98 |  | 0.05 |
| SD025 < SD100 | 0.20 | 22.00 | 0.98 |  | 0.09 |
| SD025 > SD100 | 0.30 | 24.00 | 1.00 |  | 0.02 |
| SD025 < SD100 | 0.40 | 22.00 | 0.98 |  | 0.09 |
| SD025 < SD100 | 0.50 | 9.00 | 0.14 |  | 0.53 |
| SD025 > SD100 | 0.60 | 41.00 | 0.13 |  | 0.56 |

The alternative hypotheses are determined by comparing the median growth rates of the two conditions at the given salinity stress. The p values are those after the false discovery rate correction, and the checkmarks in the column of Significance represent the p values after the correction are smaller than 0.05.

**Table S3:** Summary of one-sided Wilcoxon rank sum tests on the growth rates of the evolved strains in 1M NaCl.

| Pair and alternative hypothesis | $W$ | P value | Significance | Effect size ($r$) |
| --- | --- | --- | --- | --- |
| Control < Linear | 0.00 | 0.01 | ✓ | 0.76 |
| Control < SD025 | 0.00 | 0.01 | ✓ | 0.76 |
| Control < SD100 | 0.00 | 0.01 | ✓ | 0.76 |
| Linear > D025 | 30.00 | 0.27 |  | 0.19 |
| Linear < SD100 | 3.00 | 0.01 | ✓ | 0.73 |
| SD025 < SD100 | 5.00 | 0.01 | ✓ | 0.67 |

The results of the one-sided Wilcoxon rank-sum tests are shown. The $p$ values are those after the false discovery rate correction, and the check marks in the column of Significance represent the $p$ values after the correction are smaller than 0.05.

**Figure S5:**
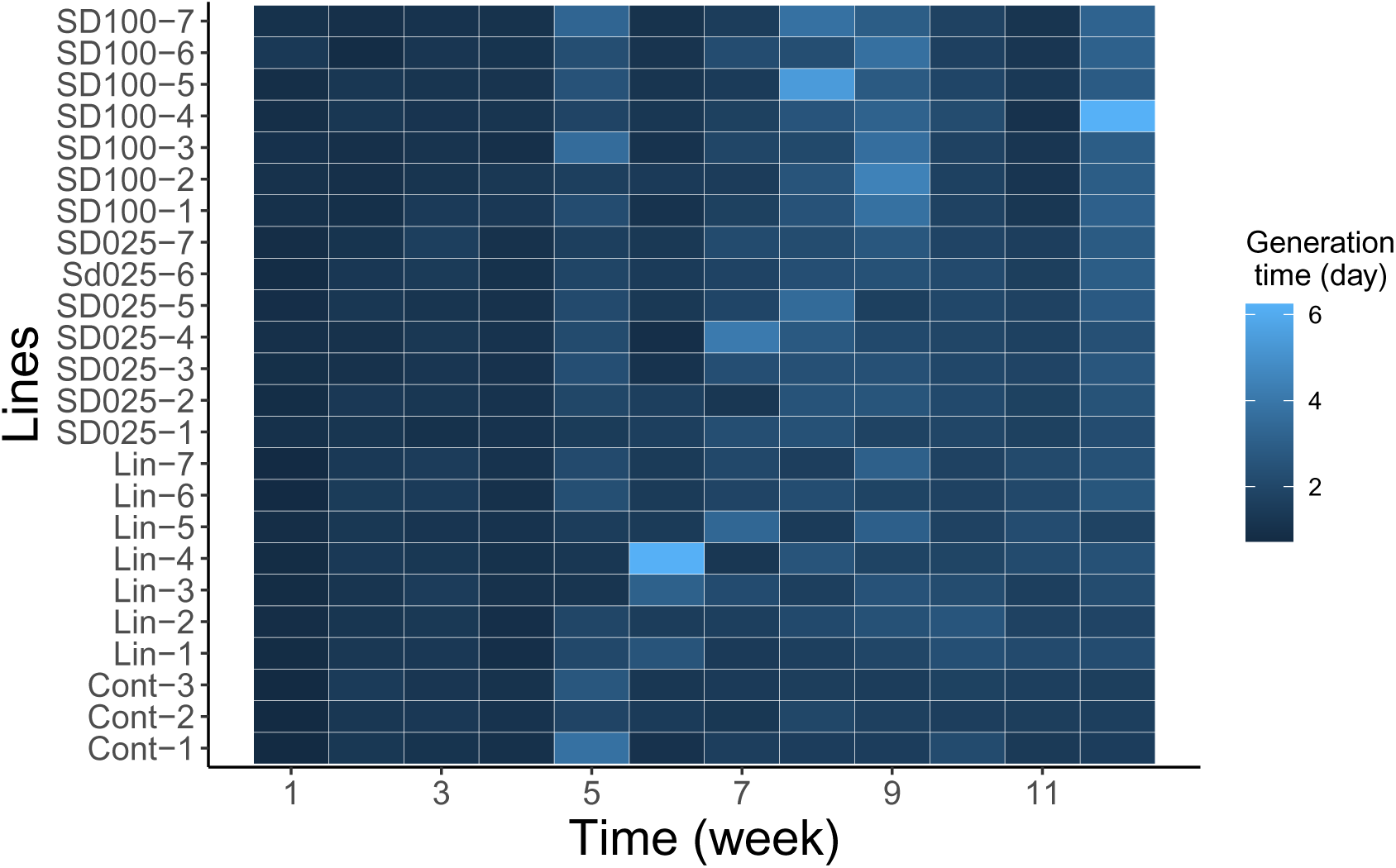
Generation time during the evolution experiment. The generation time of each strain at each week was given by 7 *× {*(log_10_ *N*_7_ *−* log_10_ *N*_0_)*/* log_10_ 2*}^−^*^1^.

**Table S4:** Details of the linear regression model in Fig S6.

|  | Estimate | Std. Error | t value | Pr(>\|t\|) |
| --- | --- | --- | --- | --- |
| (Intercept) | 0.5079 | 0.0244 | 20.80 | 0.0000 |
| NaCl | -0.7812 | 0.0740 | -10.56 | 0.0000 |

**Figure S6:**
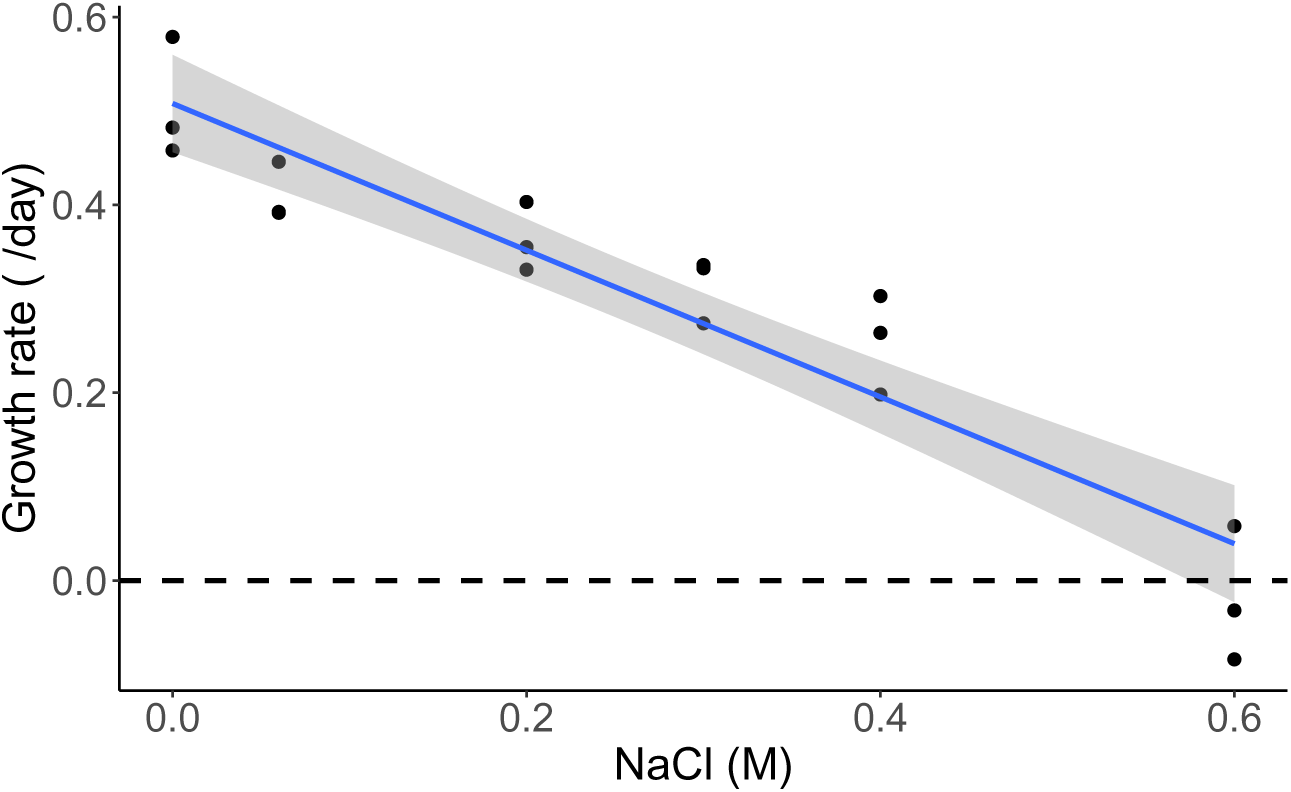
Original growth rates over the salt concentration. *C. vulgaris* was grown in 2 ml of C medium containing NaCl from 0M to 0.6M for seven days in 6-well plates. The light intensity, temperature, and the shaker’s rotation speed were identical to those in the evolution experiment explained in the main text. We have three replicates under each salt concentration. The growth rates were calculated by log(OD_750_(7)*/*OD_750_(0))*/*7, where OD_750_(*t*) the optical density at 750nm on day *t* from the culture. Because some samples show negative OD_750_(0), we added 0.001 *−* min OD_750_(*t*) to all row OD_750_(0). The optical density was measured by Multiskan SkyHigh (Thermo Fisher Scientific). The blue line represents the linear regression model, and the shaded area represents a 95% confidence interval. See Table S4 for the details of the linear regression model obtained from the lm() function in R. The dashed line represents the growth rate of 0.

**Figure S7:**
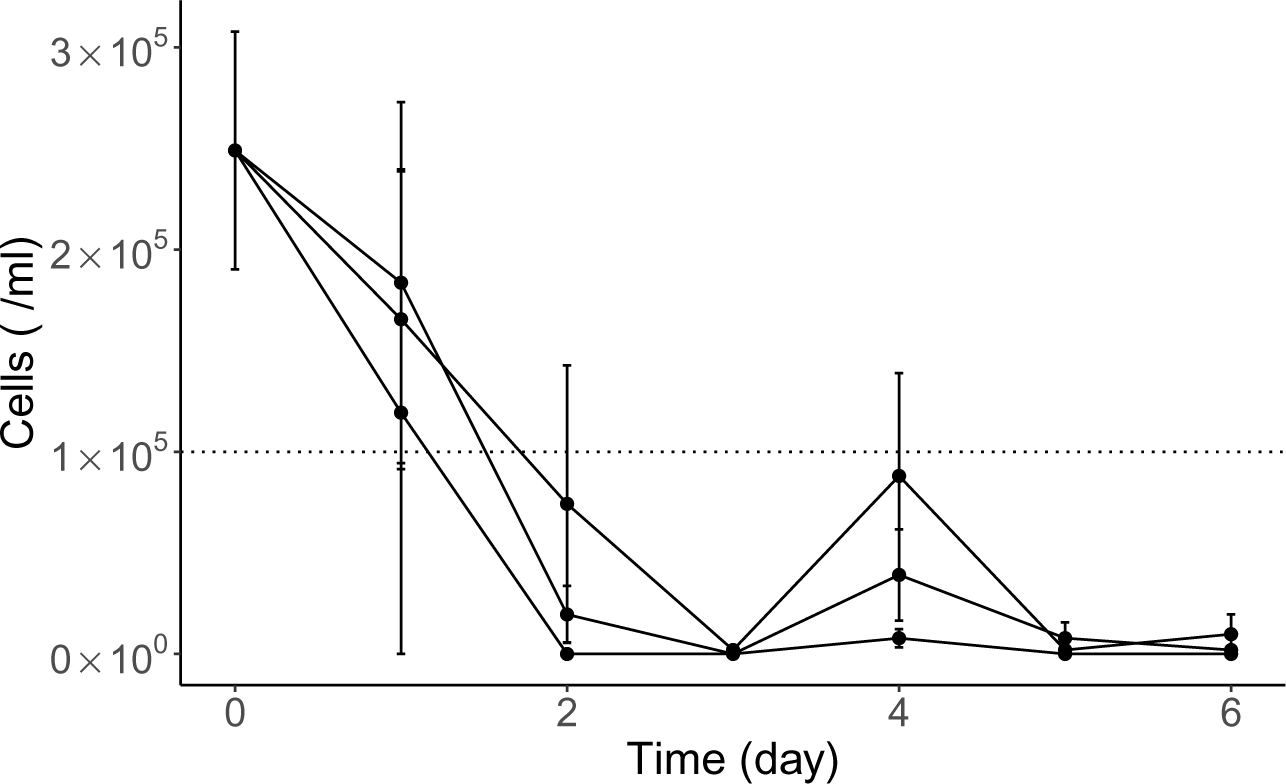
Dynamics of the original strain under 1M NaCl. *C. vulgaris* was grown in 20 ml of C medium containing 1M NaCl in an Erlenmeyer flask for six days. The light intensity, temperature, and the shaker’s rotation speed were identical to those in the evolution experiment explained in the main text. The cell density was measured by Countess II FL. The three solid lines represent the dynamics in three replicates; the dots indicate the mean of three measurements per sample, and the error bars show the standard errors. The dotted line represents the threshold that Countess II FL can accurately estimate the cell density. The algae were regarded as extinct when the cell density was lower than this threshold.

**Figure S8:**
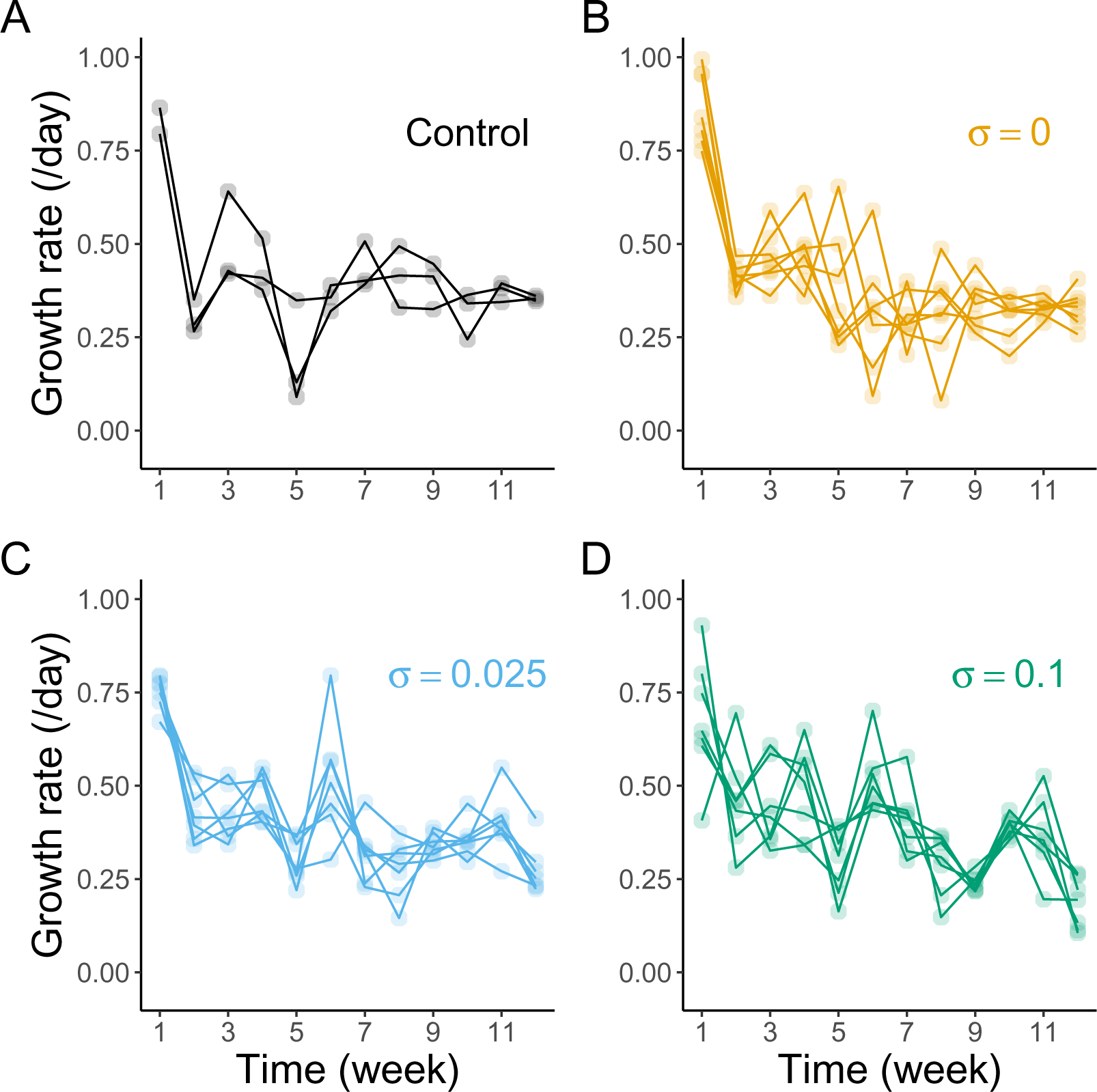
Growth rates of evolved strain over time. The growth rates in the control (A), linear (B), small-fluctuation (C), and large-fluctuation (D) conditions during the evolution experiment over time.

